# Biophysical modeling of the neural origin of EEG and MEG signals

**DOI:** 10.1101/2020.07.01.181875

**Authors:** Solveig Næss, Geir Halnes, Espen Hagen, Donald J. Hagler, Anders M. Dale, Gaute T. Einevoll, Torbjørn V. Ness

## Abstract

Electroencephalography (EEG) and magnetoencephalography (MEG) are among the most important techniques for non-invasively studying cognition and disease in the human brain. These signals are known to originate from cortical neural activity, typically described in terms of current dipoles. While the link between cortical current dipoles and EEG/MEG signals is relatively well understood, surprisingly little is known about the link between different kinds of neural activity and the current dipoles themselves. Detailed biophysical modeling has played an important role in exploring the neural origin of intracranial electric signals, like extracellular spikes and local field potentials. However, this approach has not yet been taken full advantage of in the context of exploring the neural origin of the cortical current dipoles that are causing EEG/MEG signals.

Here, we present a method for reducing arbitrary simulated neural activity to single current dipoles. We find that the method is applicable for calculating extracranial signals, but less suited for calculating intracranial electrocorticography (ECoG) signals. We demonstrate that this approach can serve as a powerful tool for investigating the neural origin of EEG/MEG signals. This is done through example studies of the single-neuron EEG contribution, the putative EEG contribution from calcium spikes, and from calculating EEG signals from large-scale neural network simulations. We also demonstrate how the simulated current dipoles can be used directly in combination with detailed head models, allowing for simulated EEG signals with an unprecedented level of biophysical details.

In conclusion, this paper presents a framework for biophysically detailed modeling of EEG and MEG signals, which can be used to better our understanding of non-inasively measured neural activity in humans.

**Graphical abstract:** 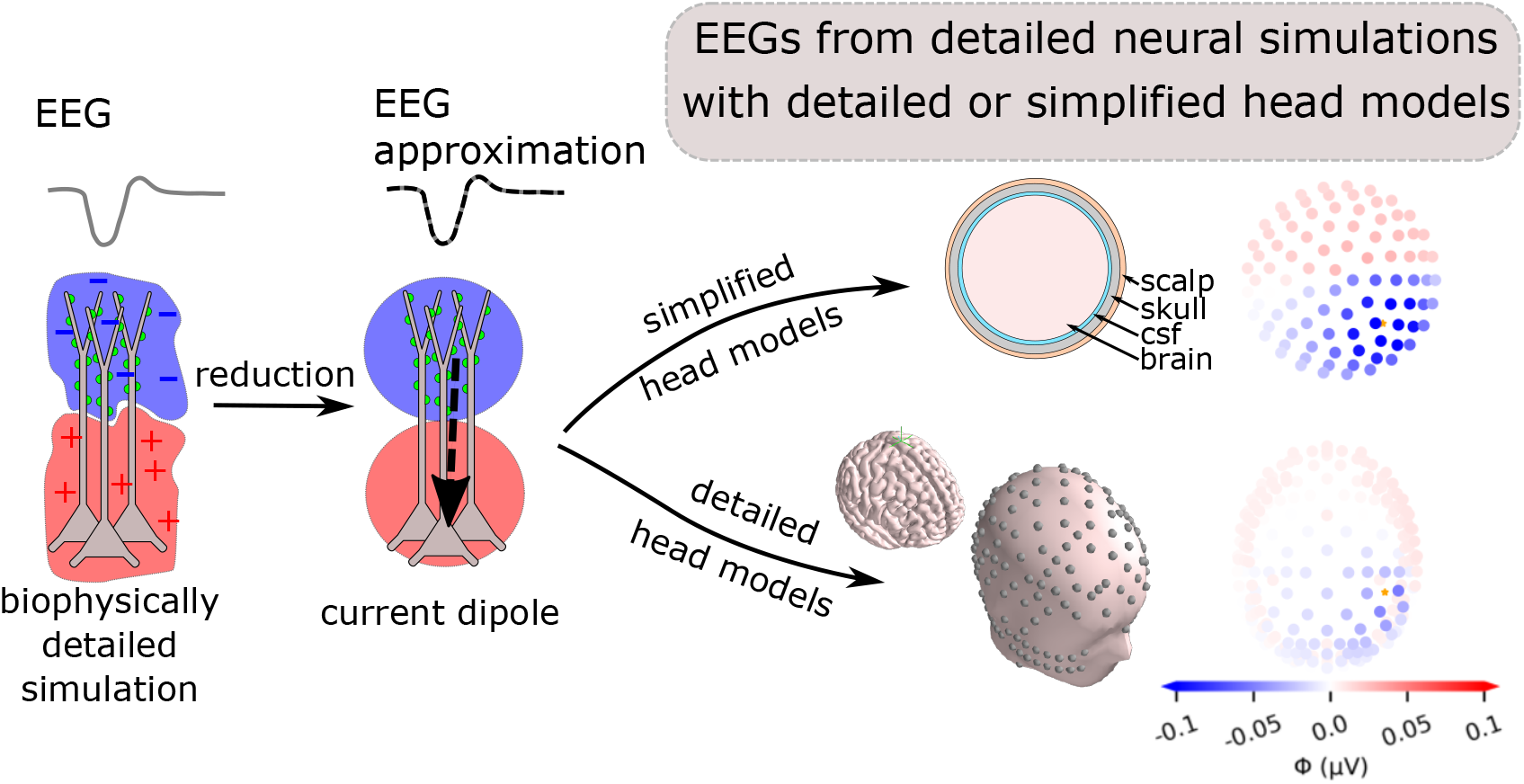

**Highlights:**

- Current dipoles are computed from biophysically detailed simulated neuron and network activity
- Extracted current dipoles allow for accurate computation of EEG and MEG signals in simplified and detailed head models
- Current-diplole approximation generally not suitable for accurate calculations of ECoG signals
- Method provides biophysics-based link between detailed neural activity and systems-level signals

## 1. Introduction

Electroencephalography (EEG) is one of the most important non-invasive methods for studying human cognitive function and diagnosing brain diseases [Cohen, 2017; Pesaran et al., 2018]. Yet, we know surprisingly little about the neural origin of these electric scalp potentials [Cohen, 2017]: On the one hand, we have a relatively good understanding of the biophysics of EEGs, in knowing that these signals originate from cortical current dipoles, and having a well-defined framework for linking such cortical dipoles to electric scalp potentials [Nunez and Srinivasan, 2006]. This has been taken advantage of for a long time in source localization, by inverse modeling of the underlying cortical current dipoles from EEG data. On the other hand, even though these cortical dipoles are assumed to mainly originate from large numbers of synaptic input to cortical pyramidal cell populations [Nunez and Srinivasan, 2006; Lopes da Silva, 2013; Pesaran et al., 2018; Ilmoniemi and Sarvas, 2019], the precise link between cortical dipoles and the underlying neural activity has remained unclear. In other words, we know very little about exactly which types of neural activity that cause even the most well-studied characteristics of the EEG signal, such as different types of oscillations (e.g., alpha, beta, and gamma waves) and stereotyped EEG shapes in response to sensory stimuli (event-related potentials, ERPs) [Cohen, 2017]. Importantly, these different EEG characteristics are affected in predictable ways by various brain conditions, such as sleep and attention [Klimesch et al., 1998; Palva and Palva, 2011; Siegel et al., 2012], and by brain disorders including epilepsy and schizophrenia [Niedermeyer, 2003; Light and Näätänen, 2013; Freestone et al., 2015; Mäki-Marttunen et al., 2019a]. This means that a better insight into how different types of brain activity is reflected in cortical current dipoles could help us not only in making better inverse models for source localization, but also in providing a better understanding of the mechanisms of human cortical activity and possibly curing brain diseases [Uhlirova et al., 2016; Cohen, 2017; Mäki-Marttunen et al., 2019a].

The reasons why we lack understanding of the neural origin of EEG signals are many, the main being (i) strong ethical constraints on invasive human brain measurements and (ii) the high number of neurons that contribute to the signal. However, in recent years there have been major advances in several relevant branches of neuroscience, meaning that a better understanding of the EEG signal may now be within reach [Uhlirova et al., 2016; Cohen, 2017].

To bypass challenge (i), we look to the rapid development in the technology and methods used to study neural activity in lab animals. The possibility to control and manipulate neural activity, while simultaneously recording both intracranial signals like the local field potential (LFP) [Einevoll et al., 2007; Blomquist et al., 2009] and extracranial non-invasive signals like the EEG [Bruyns-Haylett et al., 2017], can be expected to make important contributions to our understanding of non-invasive measurements of human brain activity [Lopes da Silva, 2013; Uhlirova et al., 2016; Cohen, 2017; Pesaran et al., 2018]. Furthermore, detailed biophysical modeling of neural activity has become an important tool for understanding intracranial LFP measurements [Einevoll et al., 2013a; Pesaran et al., 2018]. Given that EEG is expected to reflect the same basic process as LFP, that is, large numbers of synaptic input to geometrically aligned pyramidal cells [Nunez and Srinivasan, 2006; Pesaran et al., 2018; Buzsáki et al., 2012], it seems likely that detailed biophysical modeling can also help shed light on the neural origin of EEG signals.

As indicated in challenge (ii), EEG signals are expected to reflect the activity of much larger neural populations than the LFPs, making the simulations computationally demanding. Biophysically detailed large-scale simulations of neural networks have, however, been gaining substantial momentum in recent years, thanks to large ongoing neuroscience initiatives like Project MindScope at the Allen Institute for Brain Science, the Blue Brain Project and the EU Human Brain Project [Einevoll et al., 2019]. The possibility to calculate EEG signals from such existing and future large-scale biophysically detailed neural simulations could lead to valuable insights into the neural origin of the EEG.

Another complicating aspect of EEG modeling, is that these predictions in general require a head model to account for the widely different electrical conductivities of the brain, cerebrospinal fluid (CSF), skull and scalp [Nunez and Srinivasan, 2006; Ilmoniemi and Sarvas, 2019]. While many such head models exist, they tend to take current dipoles as input [Nunez and Srinivasan, 2006; Pesaran et al., 2018], instead of the transmembrane currents that are available from biophysical neural simulations and that form the basis for modeling LFPs [Einevoll et al., 2013b].

Here, we introduce an approach for reducing arbitrary biophysically detailed simulated neural activity to current dipoles, which represents an enormous reduction in term of model complexity when computing brain signals. We verify that the approach gives accurate results when calculating EEG signals, but less so for intracranial electrocorticography (ECoG) signals. Next, we look into how the approach can be applied for investigating the origin of EEG signals, with a particular focus on calcium spikes, before demonstrating how our methods can be applied for pre-existing large-scale network models. Finally, we show how current dipoles can be combined with detailed head models, which enables simulation of EEG signals with unprecedented biophysical detail.

Note that the clear separation between calculation of current dipoles and the corresponding EEG is equally valid for magnetoencephalography (MEG) signals. While we here focus mostly on EEG, the presented approach for calculating current dipoles from neural activity is equally valid for MEG signals, through use of an appropriate forward model [Hagen et al., 2018; Ilmoniemi and Sarvas, 2019].

## 2. Methods

Neural activity generates electric currents in the brain, which in turn create electromagnetic fields. In this section, we explain how electric brain signals can be modeled in both simple and more complex volume conductors.

### 2.1. Forward modeling of electric potentials

We assume negligible capacitive effects in the head [Pfurtscheller and Cooper, 1975; Ranta et al., 2017; Miceli et al., 2017] and that electric and magnetic signals effectively decouple. We can then apply the quasistatic approximation of Maxwell’s equations for calculating these signals [Hämäläinen et al., 1993; Nunez and Srinivasan, 2006]. In other words, for computing extracellular electric potentials, we envision the head as a 3D volume conductor, and combining Maxwell’s equations with the current conservation law, we obtain the Poisson equation for computing extracellular potentials [Griffiths, 1999]:

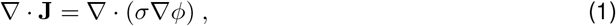

where **J** is the electric current density in extracellular space, *σ* is the extracellular conductivity and *ϕ* is the extracellular electric potential. The Poisson equation can be solved analytically for simple, symmetric head models, such as an infinitely big space and spherically symmetric models. For more complex head models, numerical methods such as the Finite Element Method (FEM) can be used [Logg et al., 2012; Vorwerk et al., 2014; Haufe et al., 2015; Seo et al., 2016; Vorwerk et al., 2019].

#### 2.1.1. Compartment-based approach

Extracellular potentials generated by transmembrane currents can be calculated with a well-founded biophysical two-step forward-modeling scheme. The first step involves *multicompartmental modeling* and incorporates the details of reconstructed neuron morphologies for calculating transmembrane currents [Sterratt et al., 2011]. In the second step, Equation (1) is solved under the assumption that the extracellular medium is an infinitely large, linear, ohmic, isotropic, homogeneous and frequency-independent volume conductor. The transmembrane currents entering and escaping the extracellular medium can be seen as current sources and sinks, and give the extracellular potential *ϕ* at the electrode location **r**,

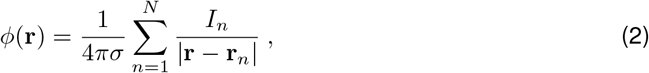

where **r**_*n*_ is the location of transmembrane current *I_n_*, *N* is the number of transmembrane currents and *σ* is the extracellular conductivity. This scheme is here referred to as the compartment-based approach, and was applied using the Python package `LFPy 2.0` running NEURON under the hood [Hagen et al., 2018; Carnevale and Hines, 2006].

#### 2.1.2. Current dipole approximation

Analogous to how electric charges can create charge multipoles, a combination of current sinks and sources can set up *current* multipoles [Nunez and Srinivasan, 2006]. From electrodynamics, we know that extracellular potentials from a volume of current sinks and sources can be precisely described by expressing Equation 2 as a multipole expansion [Nunez and Srinivasan, 2006]:

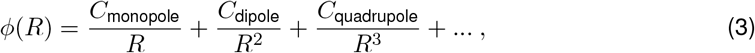

when the distance *R* from the center of the volume to the measurement point is larger than the distance from the volume center to the most peripheral source [Jackson, 1998]. In neural tissue, the current monopole contribution is zero due to current conservation, since the transmembrane currents sum to zero at all times [Koch, 1999; Pettersen et al., 2012]. Further, the quadrupole, octopole and higher order terms are negligible compared to the current dipole contribution when *R* is sufficiently large. In this case, the extracellular potential from a neuron model can be estimated with the second term of the current multipole expansion; an approximation known as the *current dipole approximation* [Pettersen and Einevoll, 2008; Pettersen et al., 2014; Nunez and Srinivasan, 2006]:

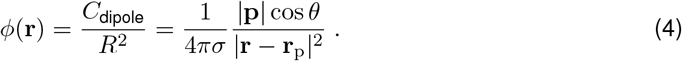

Here, **p** is the current dipole moment in a medium with conductivity *σ*, *R* = |**R**| = |**r** − **r**_p_| is the distance between the current dipole moment at **r**_p_ and the electrode location **r**, and *θ* denotes the angle between **p** and **R**. The current dipole moment **p** can be calculated from an axial current *I* inside a neuron and the distance vector **d** traveled by the axial current: **p** = *I***d**, analogous to a charge dipole moment. The current dipole approximation is applicable in the far-field limit, that is when *R* is much larger than the dipole length *d* = |**d**| [Nunez and Srinivasan, 2006].

##### Multi-dipole approach

From some multicompartmental neuron simulations (Figure 1- 3), we computed multiple current dipole moments, i.e., one for each axial current flowing between neighboring compartments in the neuron:

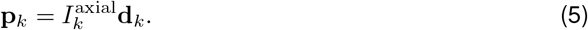

Here, 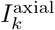 is an axial current traveling along distance vector **d**_*k*_, resulting in a current dipole moment **p**_*k*_. By inserting all the current dipole moments from a neuron simulation into the current dipole approximation (Equation 4), we get a good estimate of the extracellular potential at any electrode location where the distance between the electrode and the nearest dipole is sufficiently large [Nunez and Srinivasan, 2006]. Note that the length of each (multi-)dipole is equal to half the length of its corresponding neuronal compartment. The calculation of multi-dipoles from simulated neural activity was implemented in LFPy 2.0, and can be used through the function `Cell.get_rnulti_current_dipole_moments` [Hagen et al., 2018].

**Figure 1:**
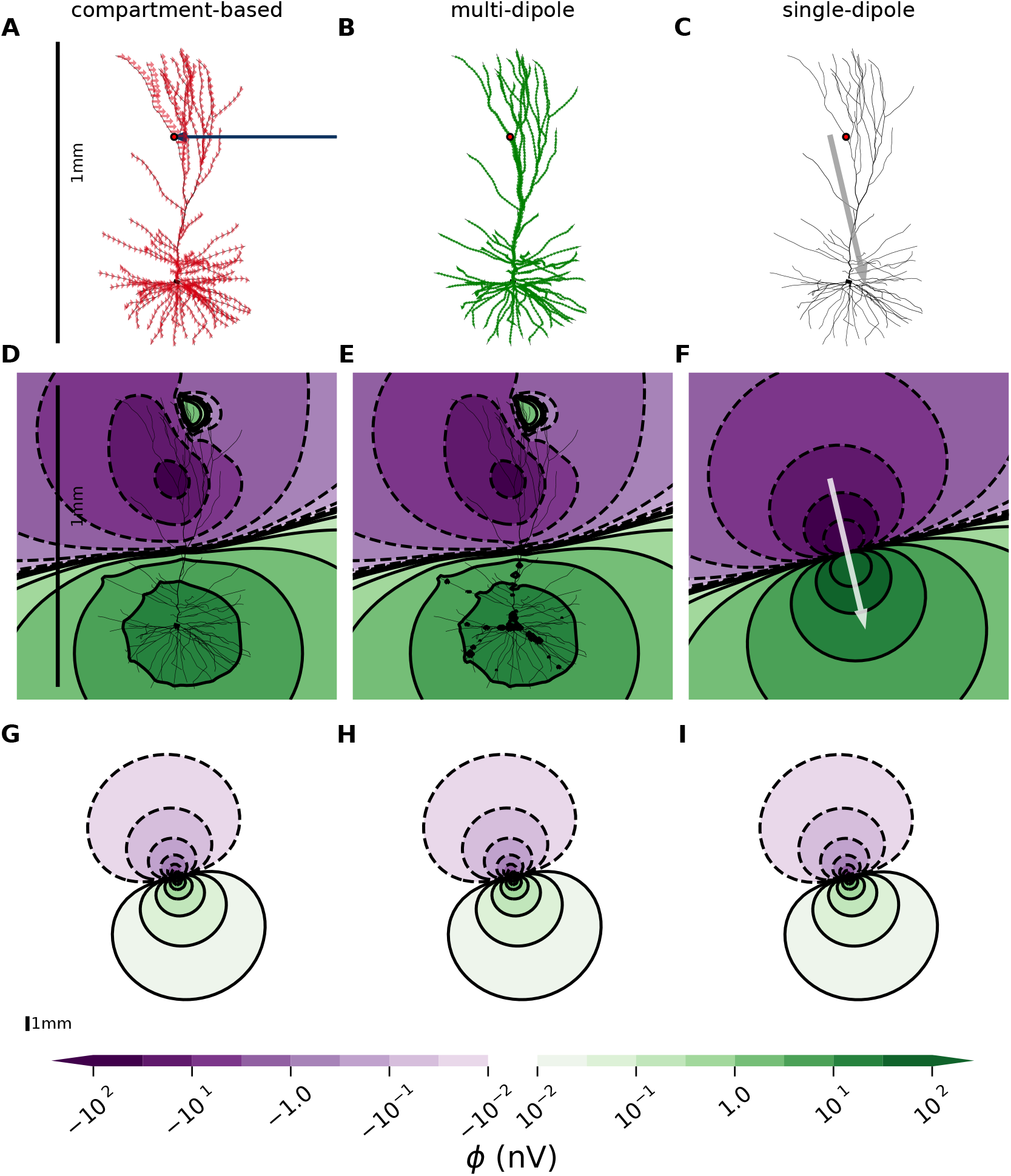
Extracellular potentials become dipolar in the far field limit. **A**: Passive l ayer-2/3 pyramidal cell from human [Eyal et al., 2016] with an excitatory, conductance-based, two-exponential synapse placed on apical dendrite (red dot), see Methods (2.3) for parameters. The resulting transmembrane currents for each compartment are shown as a blue arrow (input current) and red arrows (return currents). **B**: Green arrows represent the multiple current dipole moments between neighboring neural compartments. **C**: Gray arrow illustrates the total current dipole moment, that is, the vector sum of the dipoles in B. **D-F**: Extracellular potential in immediate proximity of the neuron, computed with the compartment-based approach, multi-dipole approach and single-dipole approximation, respectively. Note that the multi-dipole results differ slightly from the compartment-based approach when the distance from the measurement point to the nearest current dipole moment is short compared to the dipole length. **G-I**: Same as D-F, but at a larger spatial scale (zoomed out). See 1 mm scalebar in panel A, D and G. The colorbar is shared for panels D-I.

**Figure 2:**
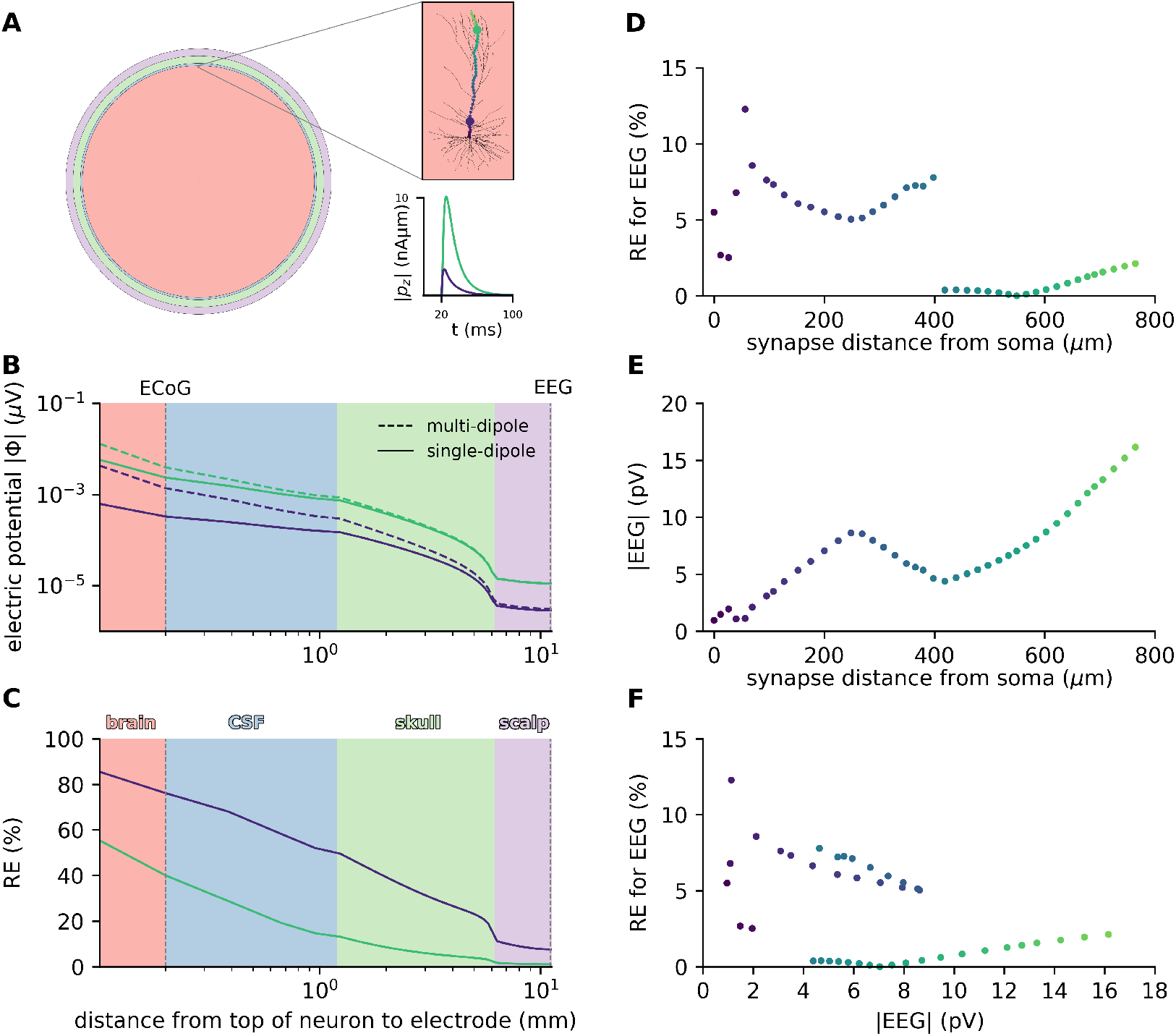
Single-dipole approximation is justified for EEG but not ECoG signals. **A**: Illustration of four-sphere head model, where the pink, blue, green and purple spherical shells represent the brain, CSF, skull and scalp respectively, see Table 1. The pink inset shows the human layer-2/3 neuron [Eyal et al., 2016] located in the brain, 78.9 mm above head center. 41 simulations lasting 100 ms with a single synaptic input after 20 ms to cell with passive ion channels only, were performed for varying input locations, see colored dots. The *z*-component of the resulting current dipole moments for two synaptic input locations (large colored dots) are shown in inset below as functions of time. The results presented in this figure are computed at the simulation time points producing the largest current dipole moment for each synaptic input location. **B**: Magnitude of extracellular potential |*ϕ*| as function of distance from the top of the neuron, shown for two simulations with synaptic input locations marked by large colored dots in upper inset of A. In each simulation, we consider the time point with the largest current dipole moment. Dashed lines show extracellular potentials computed with multi-dipole, and full lines show single-dipole calculations. **C**: Relative error RE at EEG location comparing the single-dipole model to the multi-dipole model, as function of distance from top of neuron to measurement point. **D**: Relative error RE showing how single-dipole model deviates from multi-dipole model EEG calculations, as function of distance from soma to synapse location. **E**: Magnitude of EEG signal, |EEG|, as function of distance from soma to synaptic input location. **F**: Relative error, RE, showing how EEG calculations performed with the single-dipole approximation deviates from multi-dipole approach as a function of amplitude of the EEG signal, |EEG|.

**Figure 3:**
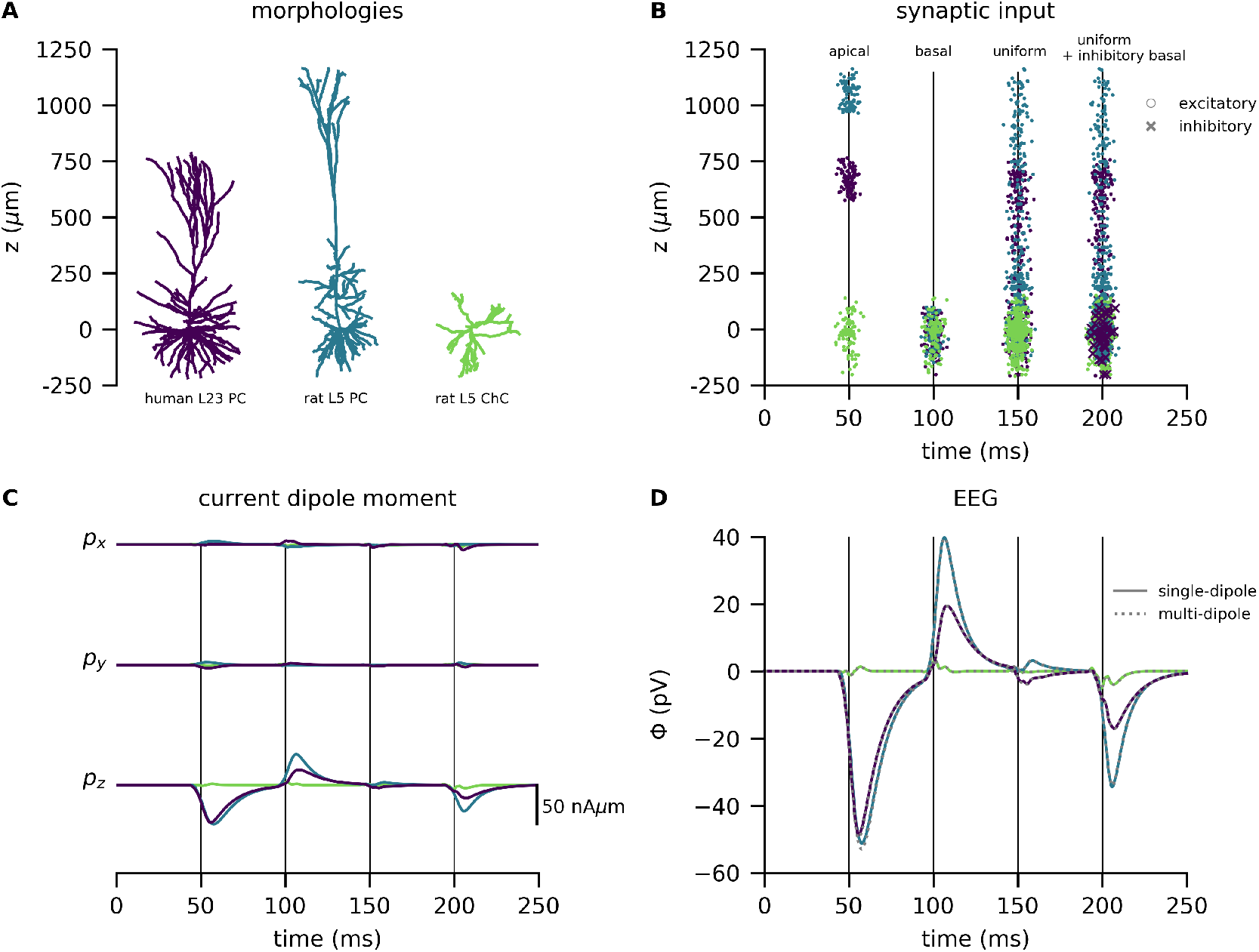
EEG signals and current dipole moment from three different cell types with various synaptic input. **A**: The morphologies of a human L2/3 pyramidal cell (purple; Eyal et al. [2016]), a rat L5 pyramidal cell (blue; Hay et al. [2011]), and a rat L5 interneuron (green; Markram et al. [2015]). The remaining panels display data connected to each cell type, see cell-specific colors. **B**: Each dot represents an excitatory synaptic input at a specific time (x-axis) at a specific height of the neuron (z-axis, corresponding to panel A) for a specific cell type (color). The crosses mark inhibitory synaptic input. The four input bulks represent 1) 100 apical excitatory synapses, 2) 100 basal excitatory input, 3) 400 homogeneously spread-out excitatory synapses and 4) 400 homogeneously spread-out excitatory synapse and inhibitory basal synapses. The synaptic weights sum to 0.01 *μ*S for all sets of excitatory/ inhibitory synapses in each wave (see sec. 2.3 for details). For the interneuron, which doesn’t have typical “apical” or “basal” zones, the synapses were spread out all over the morphology for all input types. **C**: The *x*-, *y*- and *z*-components of the current dipole moment **p** for the three different cell types. **D**: EEG signals, *ϕ* from the three cell types computed with the four-sphere model.

##### Single-dipole approximation

From each multicompartmental neuron simulation, we computed one single current dipole moment. This can either be done by summing up the multiple current dipole moments,

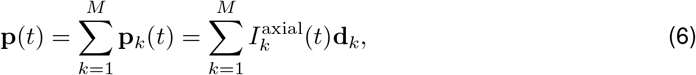

where *M* is the number of axial currents, or equivalently from a position-weighted sum of all the transmembrane currents [Lindén et al., 2010; Hagen et al., 2018]:

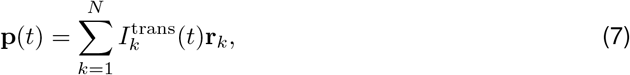

where *N* is the number of compartments in the multicompartmental neuron model and **r**_*k*_ is the position of transmembrane current 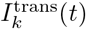. For calculating EEG signals a location for the current dipole must be chosen, and unless otherwise specified we positioned the dipole halfway between the position of the soma and the position of the synaptic input (for multiple synaptic inputs, we used the average position of the synaptic inputs). Note, however, that the large distance from the neuron to the EEG electrode (~ 10 mm) implies that the EEG signal is relatively insensitive to small changes in the dipole location within cortex. The calculation of current-dipole moments from simulated neural activity was implemented in LFPy 2.0, and can be used through `Cell.current_dipole_moment` [Hagen et al., 2018].

### 2.2. Head models

Electric potentials will be affected by the geometries and conductivities of the various parts of the head [Nunez and Srinivasan, 2006], which is especially important for electrode locations outside of the brain. This can be incorporated into our extracellular potential calculations by applying simplified or complex head models.

#### 2.2.1. Four-sphere head model

The four-sphere head model is a simple analytical model consisting of four concentric shells representing brain tissue, cerebrospinal fluid (CSF), skull and scalp, where the conductivity can be set individually for each shell [Srinivasan et al., 1998; Nunez and Srinivasan, 2006], see Table 1 for parameters used in this paper. The model solution is given in Næss et al. [2017] and is found by solving the Poisson equation subject to boundary conditions ensuring continuity of current and electric potentials over the boundaries, as well as no current escaping the outer shell. This model is based on the current dipole approximation.

**Table 1:** Radii and electrical conductivities used in the four-sphere model. The radius of each spherical shell in the four-sphere model, with *σ* denoting the respective electrical conductivities.

| | Radius (cm) | $\sigma$ (S/m) |
| --- | --- | --- |
| Brain | 7.9 | 0.3 |
| CSF | 8.0 | 1.5 |
| Skull | 8.5 | 0.015 |
| Scalp | 9.0 | 0.3 |

#### 2.2.2. New York Head model

The New York Head model is a detailed head model based on high-resolution, anatomical MRI-data from 152 adult heads [Huang and Parra, 2015]. The model was constructed by taking advantage of the reciprocity theorem, stating that the position of the electrode and the dipolar source can be switched without affecting the measured potential [Rush and Driscoll, 1969]. This means, that virtually injecting current at the locations of the EEG electrodes and using the finite element method [Logg et al., 2012] to compute the resulting potential anywhere in the brain, gives the link between current dipoles in the brain and the resulting EEG signals [Malmivuo and Plonsey, 1995; Ziegler et al., 2014; Huang et al., 2016; Dmochowski et al., 2017]. This link was captured in a matrix known as the *lead field* **L** [Nunez and Srinivasan, 2006]:

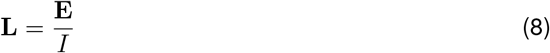

Here, *I* is the injected current at the electrode locations and **E** is the resulting electric field in the brain. The lead field matrix gives us the precise link between a current dipole moment **p** in the brain and the resulting EEG signals **Φ** [Nunez and Srinivasan, 2006]:

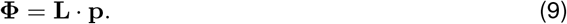

We applied the New York Head model by downloading the lead field **L** from parralab.org/nyhead/. The units incorporated in the lead field matrix was not immediately obvious. However, from Dmochowski et al. [2017]; Huang et al. [2013] it appears that an injected current *I* of 1 mA gives an electric potential *E* in V/m, meaning that a current dipole moment **p** in the unit of mAm gives EEG signals in the unit of V.

### 2.3. Simulation of neural activity

All neuron simulations were performed using the python package `LFPy`, running NEURON under the hood [Hagen et al., 2018]. For investigations of single-cell contributions to extracellular potentials, we applied three different morphologically reconstructed cell models: The human layer-2/3 pyramidal cell from Eyal et al. [2018], the layer-5 pyramidal cell from rat cortex constructed by Hay et al. [2011] and a rat layer-5 chandelier cell; an interneuron model developed by Markram et al. [2015].

The pyramidal cell models were downloaded from senselab.med.yale.edu/modeldb/, with accession numbers 238347 (2013_03_06_cell03_789_H41_03) and 139653 (cell1) respectively, while we found the interneuron at the Neocortical Microcircuit Collaboration Portal (bbp.epfl.ch/nmc-portal/microcircuit) under layer-5, Chandelier Cell (ChC), continuous Non-accomodating (cNAC), (rp110201_L_idA_-_Scale_x1.000_y0.975_z1.000_-_Clone_3).

For all simulations with passive ion channels only (Fig. 1-3), we used the following cell parameters: membrane resistance of 30000 Ωcm^2^, axial resistance of 150 Ωcm [Mainen and Sejnowski, 1996] and a membrane capacitance of 1 *μ*F*/*cm^2^ [Gentet et al., 2000; Sterratt et al., 2011]. When active mechanisms were included in the simulations (Fig. 4), all cell properties were incorporated as described in the specific cell’s documentation.

**Figure 4:**
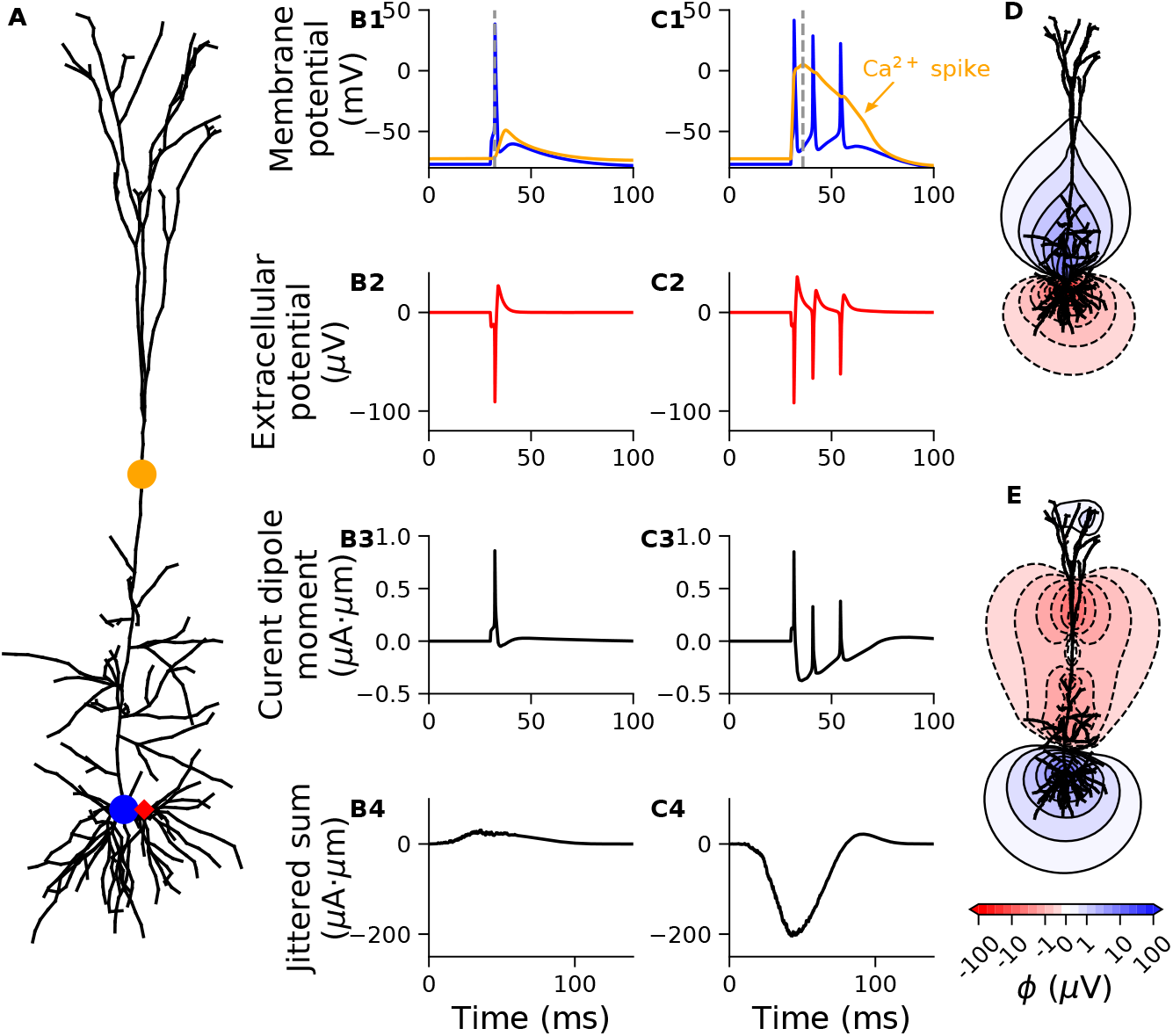
Current dipole moment expose dendritic calcium spikes. **A**: Layer-5 cortical pyramidal cell model from rat [Hay et al., 2011], receiving either a single excitatory synaptic input to the soma evoking a single somatic action potential (blue dot, results in **B1-4**), or in addition an excitatory synaptic input to the apical dendrite, evoking a dendritic calcium spike and two additional somatic spikes (orange dot, results in **C1-4**). **B1, C1**: Membrane potential at the two positions indicated in **A**. **B2, C2**: Extracellular potential 30 μm away from the soma (red diamond in **A**), assuming for illustration an infinite homogeneous extracellular medium. **B3, C3**: Single-cell current dipole moment. **B4, C4**: Sum of 1000 instances of the single-cell current dipole moment (from **B3, C3**), that has been randomly shifted in time with a normally distributed shift with a standard deviation of 10 ms. **D:** Contour lines of extracellular potential around neuron at a snapshot in time during the somatic spike in **B1** (*t*=32.2 ms; time marked by dashed line). **E:** Contour lines of extracellular potential around neuron at a snapshot in time during the calcium spike in **C1** (*t*=36.0 ms; time marked by dashed line). The synaptic weight was 0.07 and 0.15 *μ*S for the somatic and apical input location, respectively.

Neural simulations shown in Fig. 1-4 received synaptic input modeled as conductance-based, two-exponential synapses (`Exp2Syn` in `NEURON`). The rise time constant was set to 1 ms and the decay time constant was 3 ms, synaptic reversal potential was 0 mV and the synaptic weight was set to 0.002 *μ*S, unless otherwise specified.

For modeling of population activity (Figure 5, 6), we used the so-called hybrid scheme recently proposed by Hagen et al. [2016]. The simulation was unmodified from their presented results with transient thalamocortical input (their Fig. 1 and 7), except that all single-cell current dipole moments were recorded, and the EEG signals calculated.

**Figure 5:**
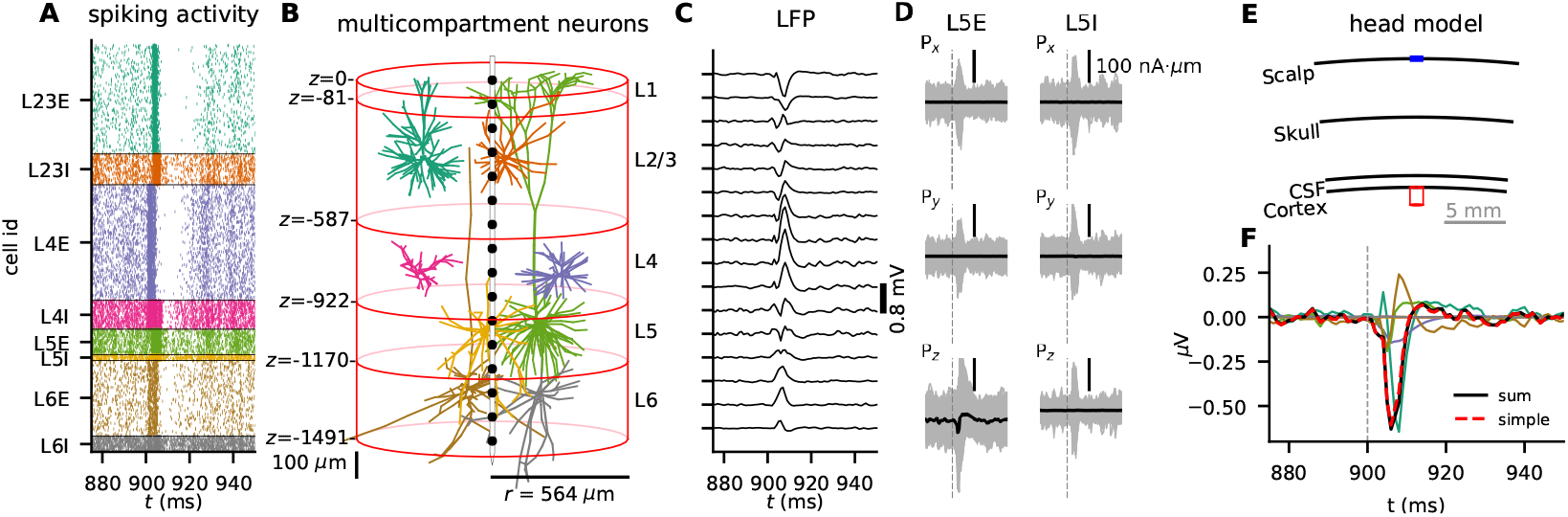
Large-scale neural simulations can be used to probe biophysical origin of EEG signals. **A**: Stimulus-evoked spiking activity from thalamic input (time *t* = 900 ms, denoted by thin vertical line) in the cortical microcircuit model from Potjans and Diesmann [2014]. Dots indicate spike times of individual neurons, and populations are represented in different colors (I=inhibitory, E=excitatory). **B**: Multicompartment model neurons used to produce the measurable signals, with colors corresponding to panel **A**, showing one example morphology per population. Layer boundaries are marked at depths relative to cortical surface, z = 0. A laminar recording electrode with 16 contacts separated by 100 *μ*m (black dots) is positioned in the center of the population. **C**: LFPs calculated at depths corresponding to black dots in **B**. **D**: For the two L5 populations (L5I and L5E), the three components of the current dipole moment is shown for all individual cells (gray), together with the population average (black). **E**: Illustration of the four-sphere head model, with the red column corresponding the the outline of the population in panel **B**. **F**: The EEG signal from each population found by summing the single-cell EEG contribution of all individual cells within each population (different colors, same color scheme as in **A**,**B**), together with the total summed EEG signal (black). The simplified EEG signal was found by first summing the *z*-component of the current dipole moments for all pyramidal cells, that is L2/3E, L5E and L6E, and calculating the EEG from this single current dipole (red dashed).

**Figure 6:**
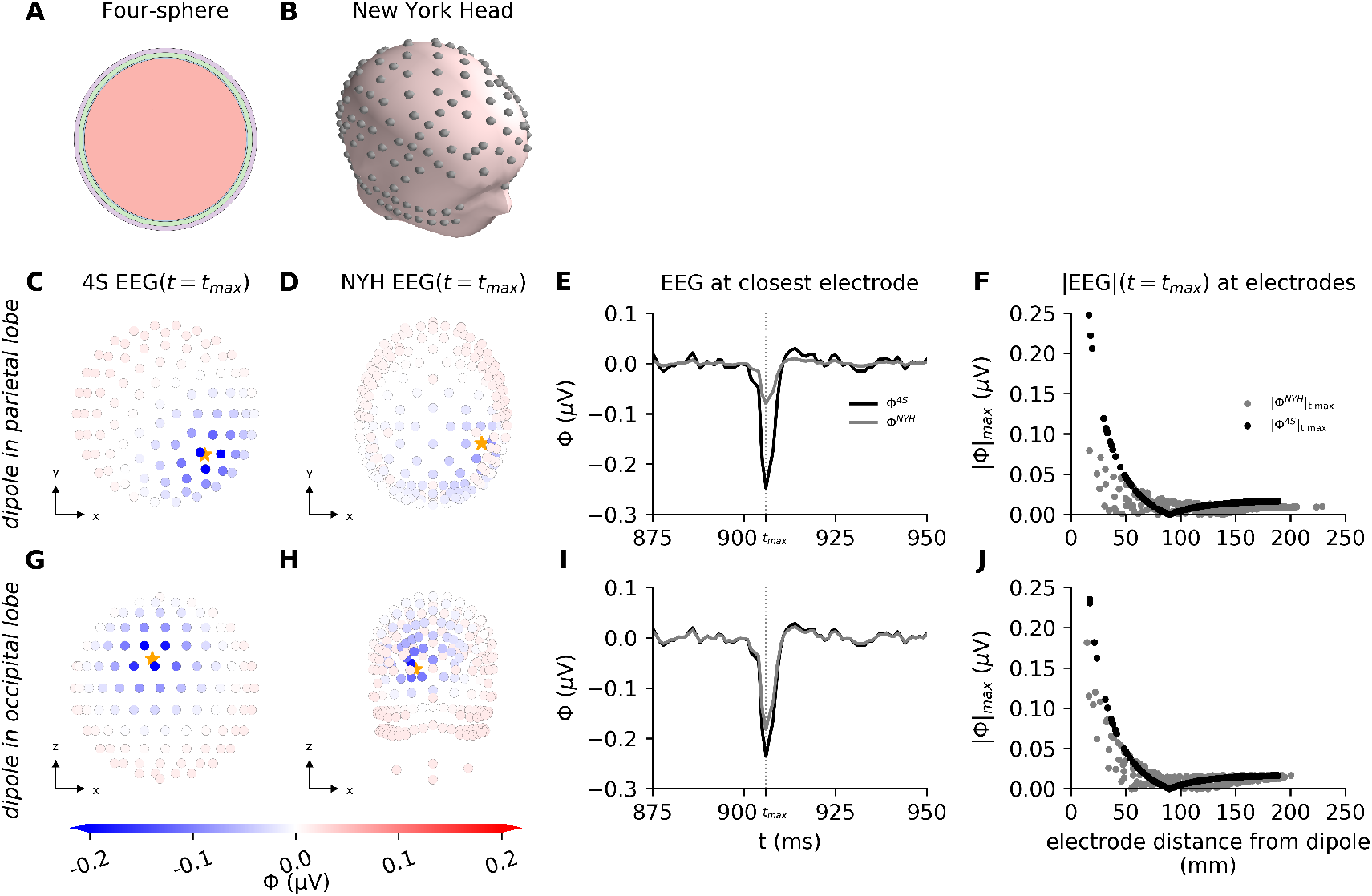
EEG signals from cortical column network can be modeled with the four-sphere model and the New York Head. EEG signals from population dipole resulting from waves of synaptic input to the cortical microcircuit model from Potjans and Diesmann [2014]. Population dipole was placed in two different locations: parietal lobe (C,D,E,F) and occipital lobe (G,H,I,J). **A**: The four-sphere model consisting of four concentrical shells: brain tissue, CSF, skull and scalp. **B**: The New York Head model. **C, D**: EEG signals (*ϕ*) on scalp surface electrodes, seen from above, showing time point of the strongest current dipole moment |**p**| of the population simulation. Dipole is placed in the parietal lobe and location is marked by orange star, having the following coordinates in the NYH model: (55, −49, 57) mm. EEG signals were computed with the New York Head model (C) and the four-sphere head model (D). **E**: EEG trace computed with the four-sphere model (black) and the New York Head model (gray) on closest scalp surface electrode: i.e. the electrode with the shortest distance to the current dipole moment location. Dipole is placed in parietal lobe (distance is 16.13 mm for the four-sphere model and 16.76 mm for NYH). **F**: Absolute value of EEG signals from panel C,D generated by dipole in parietal lobe, plotted as function of distance from dipole to measurement electrode. **G,H**: Equivalent to panel C,D, however, dipole is placed in occipital lobe, and electrodes are seen from the back of the head. Dipole coordinates for NYH model: (−24.3, −105.4, −1.2) mm. **I**: EEG trace from dipole in occipital lobe computed on closest electrode (distance is 16.90 mm for the four-sphere model and 14.64 mm for NYH), equivalent to panel E. **J**: Equivalent to panel F, with EEG signals from panel G,H generated by dipole in occipital lobe.

### 2.4. Code availability

Simulation code to reproduce all figures in this paper is freely available from https://github.com/solveignaess/EEG.git

## 3. Results

We introduce an approach for modeling electroencephalography (EEG) and magnetoencephalography (MEG) signals from detailed biophysical multicompartment cell models. The approach involves two steps: First, current dipole moments are extracted from activity in neurons or networks. Second, the extracted current dipoles are used as sources in established forward models. Here we only demonstrate the approach by computing EEG signals, but the current dipoles are equally applicable for computing MEG signals using the appropriate magnetic-field forward models [Hämäläinen et al., 1993; Hagen et al., 2018; Ilmoniemi and Sarvas, 2019]. For illustration, we first consider EEG signals stemming from single synaptic input onto single neurons in an infinite homogeneous head model, before moving on to a simple, generic head model. Finally, we study EEGs from large-scale simulated network activity, also applying a detailed head model.

### 3.1. At sufficiently large distances, extracellular potentials become dipolar

When modeling electric potentials within the brain, we can apply the well-established compartment-based approach assuming a homogeneous volume conductor (section 2.1.1) [Einevoll et al., 2013a; Holt and Koch, 1999]. However, this assumption is no longer valid when it comes to modeling EEG signals on the scalp, which calls for an inhomogeneous head model [Ilmoniemi and Sarvas, 2019]. Such head models typically take current dipoles as input, as opposed to individual current sinks/sources, and must be based on the current dipole approximation [Nunez and Srinivasan, 2006]. Here, we introduce an approach for computing current dipoles from arbitrary simulated neural activity, and compare current-based and dipole-based modeling of electric potentials generated by a single cell receiving excitatory synaptic input. Excitatory synaptic input initiates a negative current at the synapse location, since positive ions flow into the cell. Due to current conservation [Koch, 1999], this negative current is exactly balanced by spatially distributed positive currents along the cellular membrane, as illustrated in Fig. 1**A** for a single apical excitatory synaptic input to a passive human cortical layer-2/3 pyramidal cell model [Eyal et al., 2016]. See Methods 2.3 for simulation details. In the standard procedure for modeling extracellular potentials, here referred to as the *compartment-based approach*, the transmembrane current in each cellular compartment corresponds to a point current source/sink. Another strategy is to consider the axial current of each cellular compartment as a small current dipole (see Equation (6)), which we refer to as the *multi-dipole approach* (Fig. **1B**). By vector summation of all these dipoles into one single dipole at a specific position, we obtain the *single-dipole approximation* (Fig. **1C**). For the sake of comparing these modeling approaches, we have assumed that the cell is positioned in an infinite homogeneous electric medium. Very close to the neuron, the extracellular potential will strongly depend on the exact distribution of transmembrane currents across the cellular morphology and will, therefore, typically not take a purely dipolar shape (Fig. 1**D,E** versus **F**). However, since the dipole contribution will dominate when we are further away from the current sources (see Equation 3), the extracellular potential becomes more and more dipolar with increasing distance from the cell [Lindén et al., 2010]. This implies that for the purpose of calculating extracellular potentials far away from the cell, the single-dipole approximation might be well justified (Fig. 1**G-I**). Note that there can be small differences between the results from the compartment-based and the multi-dipole approaches for electrode locations in the immediate vicinity of the current sources, due to the approximations inherent in using the current dipole model (Fig. 1D versus Fig. 1E).

### 3.2. Single-dipole approximation is justified for EEG, but not ECoG signals

In order to test the applicability of the single-dipole approximation for calculating ECoG and EEG signals, we applied the four-sphere head model [Næss et al., 2017; Hagen et al., 2018, 2019]. Since the four-sphere head model takes current dipoles as input, the multi-dipole approach was used as benchmark: an assumption that should be well justified for the cell-to-electrode distances considered, see section 3.1.

For different locations of a single excitatory synaptic input to a human cortical layer-2/3 pyramidal cell model [Eyal et al., 2016] (Fig. **2A**), we calculated the electric potential at point-electrode positions spanning from 100 *μ*m above the top of the cell, to the surface of the head, using both the multi-dipole approach and the single-dipole approximation (Fig. **2B**). In the simulations shown, we used conductance-based synapses and included only passive membrane conductances, but we confirmed that using current-based synapses or a fully active cell model gave very similar results.

The electric potential decreased steeply with distance when crossing the different layers of the head model, most strongly across the low-conducting skull (Fig. **2B**). For all synaptic input locations, we observed that the electric potential calculated with the single-dipole approximation markedly deviated from the multi-dipole approach directly above the neuron, but the difference strongly decreased with distance from the neuron (Fig. **2B**, full versus dashed lines for two selected synapse locations). We quantified the model dissimilarities by looking at the relative error at the timepoint of the maximum current dipole moment, and for a chosen distal synaptic input the relative error was 40.0% and 1.06% at the position of the ECoG and EEG electrodes respectively (Fig. **2C**, green line). For a specific proximal synaptic input we observed a relative error of 76.1% a t the ECoG position, and 7.61% at the EEG position (Fig. **2C**, purple line). Inserting a single strong synaptic current (synaptic weight 0.05 *μ*S) into the soma of the same layer-2/3 pyramidal cell with active mechanisms [Eyal et al., 2018], resulting in a somatic spike, gave relative errors of 34.7% and 0.967% for the computed ECoG and EEG signals, respectively (results not shown). We found that calculating EEG signals with the single-dipole approximation gave relative errors peaking for synaptic locations ~ 60 and 400 μm above the soma (Fig. **2D**), but note that these synaptic input locations also gave relatively weak EEG signals (Fig. **2E**). This demonstrates that the relative error of the single-dipole approximation is negatively correlated with the amplitude of the scalp potential (Fig. **2F**). This is as expected, given that the strongest EEG signals are expected to be caused by dipole-like source/sink distributions (section 2.1.2). In summary, the single-dipole approximation can result in substantial errors at the position of the ECoG electrodes, but gives small errors at the position of the EEG electrodes for synaptic locations leading to strong EEG signals.

### 3.3. Single-dipole approximation simplifies estimate of EEG contribution

In the previous section, we showed that the single-dipole approximation was applicable for calculation of EEG signals, and in this section we demonstrate that the single-dipole approximation can substantially simplify the analysis of the biophysical origin of EEG signals.

Pyramidal cells have a preferred orientation along the depth axis of cortex (here the *z*-axis), and the direction of the current dipole moment **p** can be expected to align with this axis since radial symmetry will tend to make the orthogonal components (*p_x_*, *p_y_*) cancel at the population level [Hagen et al., 2018]. In contrast, interneurons show much less of a preferred orientation, and are therefore expected to give a negligible contribution to the EEG signal, except indirectly through synaptic inputs onto pyramidal cells [Hagen et al., 2016]. We illustrated this by applying the single-dipole approximation to three different cell types (Fig. **3A**), each receiving a large number of synaptic inputs with target regions on the cells set up to vary over time (Fig. **3B**).

For the previously used human layer-2/3 cell (Fig. **3A**, purple; Eyal et al. [2016]) receiving a volley of excitatory synaptic inputs that were restricted to the uppermost 200 μm of the cell (*t*=50 ms; Fig. **3B**, purple dots), we observed a negative deviation of *p_z_* (Fig. **3C**, purple line). For basal synaptic input (*t*=100 ms; Fig. **3B**, purple line), the polarity of *p_z_* was instead positive, but of slightly lower amplitude than for apical input, as can be expected because the large area of the somatic region will cause strong return currents in the immediate vicinity of the synaptic inputs, and therefore an overall weaker current-dipole moment.

A uniform distribution of 400 synaptic inputs across the cell membrane with area-weighted probability (*t*=150 ms; Fig. **3B**, purple line), only gave rise to small ripples in *p_z_*, due to the substantial cancellation of current dipoles of opposite polarity. It is sometimes assumed that excitatory input is relatively uniformly distributed onto pyramidal cells, while inhibitory input is more directed to the perisomatic region [Mazzoni et al., 2015; Teleńczuk et al., 2019; Skaar et al., 2020; Teleńczuk et al., 2020]. As expected, we found that this combination of uniformly distributed excitatory synaptic input and perisomatic inhibitory input gave rise to a clear negative response in *p_z_* (*t*=200 ms; Fig. **3B**, purple line), which could be part of the explanation why inhibitory synaptic input in some cases has been found to dominate the LFP [Hagen et al., 2016; Teleńczuk et al., 2017].

For a rat cortical layer-5 pyramidal cell model (Fig. **3A**, blue; Hay et al. [2011]), the resulting current dipole moment was very similar in shape, but larger in amplitude, which was expected because the longer apical dendrite will tend to give larger current dipole moments (Fig. **3C**, blue line). Lastly, we used a rat cortical layer-5 interneuron model (Fig. **3A**, green; Markram et al. [2015]), but since the dendrites of interneurons are not structured into the same distinctive zones as pyramidal cells, the synaptic input caused very small net current dipole moments.

We calculated the EEG signals with the four-sphere head model, using both the multi-dipole (Fig. **3D**, dotted lines) and the single-dipole (Fig. **3D**, solid lines) approach. To compare the approaches, we computed the relative error over time, that is, the absolute difference between the results from the two approaches, normalized by the maximum EEG magnitude computed with the multi-dipole approach. The single-dipole approach gave a maximum error of 2.2%, 3.5% and 0.34% for the human layer-5 cell, the rat layer-5 cell and the rat interneuron, respectively. Importantly, the EEG signal is essentially fully described by the *z*-component of the current dipole moment *p_z_*, that is, a single time-dependent variable. This reduction in signal description represents a massive simplification in the understanding of the biophysical origin of the EEG signal, compared to considering the transmembrane currents and position of each cellular compartment.

### 3.4. Current dipole moment expose dendritic calcium spikes

Suzuki and Larkum [2017] recently demonstrated that dendritic calcium spikes can be recorded experimentally at the cortical surface, and that the signal amplitudes can be similar to contributions from synaptic inputs. This demonstrates that active conductances may play an important role in shaping ECoG and EEG signals. Furthermore, it suggests that information about calcium dynamics might be present in such signals, and that this information could potentially be taken advantage of when studying learning mechanisms associated with dendritic calcium spikes [Suzuki and Larkum, 2017].

The previously introduced rat layer-5 cortical pyramidal cell model from Hay et al. [2011] can exhibit dendritic calcium spikes. When this cell model received a single excitatory synaptic input to the soma (Fig. **4A**, blue dot), strong enough to elicit a somatic action potential (Fig. 4**B1**, blue), a small depolarization was also visible in the apical dendrite (Fig. 4**B1**, orange). Even so, this did not initiate any dendritic calcium spike. However, when combining the same somatic synaptic input with an additional excitatory synaptic input to the apical dendrite, 400 μm away from the soma (Fig. **4A**, orange dot), we observed a dendritic calcium spike. This calcium spike did, in turn, induce two additional somatic spikes (Fig. 4**C1**). For both synaptic input strategies described above, the extracellular potential simulated 30 μm away from the soma took the shape of stereotypical extracellular action potentials: that is, a sharp negative peak followed by a broader and weaker positive peak (Fig. **4B2, C2**). Further, we observed that the slow dendritic calcium spike was not reflected in the extracellular potential close to the soma (Fig. **4C2**). We found that for the case with only a somatic spike and no calcium spike, the single-cell current dipole moment resembled the inverse of the extracellular potential (Fig. 4**B3**), while for the case with both somatic and dendritic spiking, a pronounced slow component was also present in the single-cell current dipole moment (Fig. 4**C3**). Somatic action potentials are typically not expected to contribute significantly to EEG signals (but see Teleńczuk et al. [2015]), because the very short duration of spikes with both a positive and a negative phase implies that extreme synchrony is needed for spikes to sum constructively, and spikes that are only partially overlapping tend to sum destructively. The same cannot be expected to hold for the calcium spikes, which are not only longer-lasting but also predominately cause a negative response in the current current dipole moment. To mimic a neural network scenario with multiple cells spiking at slightly different times, we calculated the sum of 1000 instances of the single-cell current dipole moment that was jittered (shifted) in time (normally distributed, standard deviation=10 ms). We found that the case with the dendritic calcium spike now had a 6.6-fold larger maximum amplitude than the case with only the somatic spike (Fig. 4 **B4** versus **C4**, max |**p**| = 30.8 μAμm and 204.2 μAμm respectively). This demonstrates that dendritic calcium spikes are much more capable of summing constructively for a population of cells, and substantiates the role of dendritic calcium spikes in affecting ECoG/EEG/MEG recordings.

The amplitude of the slow component of the current dipole moment from the calcium spike was about 0.5 μAμm (Fig. 4**C3**), and later (Sec. 3.5) we will present results from a simulated neural network where the average event-related current dipole moment of layer 5 pyramidal cells were found to be about 0.1 μAμm (Fig. **5D**, bottom left). This indicates that our results are compatible with the claim by Suzuki and Larkum [2017] that signal amplitudes from calcium spikes could be similar in amplitude to contributions from synaptic input.

We can make a very rough estimate of the number of simultaneous calcium spikes required to cause a measurable EEG response: A current dipole moment of 1 μAμm gives an EEG amplitude on the order of 10^−3^ μV (see for example Fig. 3 **C**,**D**, note different scales). Assuming that an EEG contribution must exceed ~10 μV to be detectable [Nunez and Srinivasan, 2006; Hagen et al., 2018] implies a minimum needed current dipole moment of ~10^4^ μAμm. A number of perfectly synchronous calcium spikes would each contribute with ~0.5 μAμm (Fig. **4C3**), suggesting that about 20.000 synchronous calcium spikes would be needed to cause a measurable EEG response. Further, considering that the signal amplitude decreases by about 100-fold from cortical surface to scalp (Fig. **2B**) and assuming a similar detection threshold, indicates that a few hundred simultaneous calcium spikes would be detectable by ECoG electrodes.

It might initially seem surprising that the dendritic calcium spike is so strongly reflected in the single-cell current dipole moment, given that the transmembrane currents associated with the somatic action potential are much larger than those associated with the dendritic calcium spike: the maximum amplitude of the transmembrane currents of the somatic compartment was 45.1 nA, compared to just 0.30 nA for the compartment in the apical dendrite (Fig. **4A**, blue and orange dots). However, the current dipole moment is given as the product between the amplitude of the current and the separation between the source and sink (**p** = *I***d**; Equation 6). While the currents associated with the somatic action potential will for the most part be contained within the somatic region, giving very small sink/source separations, the currents associated with the dendritic calcium spike will be distributed over a much larger part of the cell membrane. This effect can be illustrated by comparing the spatial profile of the extracellular potentials around the neuron at a snapshot in time during a somatic spike or during a calcium spike (Fig. 4 **D** versus **E**).

### 3.5. EEG from large-scale neural network simulations

So far, we have only considered EEG contributions from single cells, but real EEG signals are expected to reflect the activity of hundreds of thousands to millions of cells [Nunez and Srinivasan, 2006; Cohen, 2017]. Biophysically detailed modeling of large populations is still in its infancy [Einevoll et al., 2019] and at present typically include “only” a few tens of thousands of biophysically detailed cells [Markram et al., 2015; Billeh et al., 2020]. Networks of point neurons, on the other hand, are regularly used to simulate hundreds of thousands [Billeh et al., 2020] or even millions of cells [Senk et al., 2018; Schmidt et al., 2018], but LFP, ECoG, EEG or MEG signals can not be computed directly from point neurons [Einevoll et al., 2013a]. To investigate EEG signals generated by neuronal networks, we therefore used a hybrid scheme [Hagen et al., 2016; Senk et al., 2018; Skaar et al., 2020], where the network activity is first simulated in a highly computationally efficient manner with point neurons in NEST [Linssen et al., 2018] and the resulting spiking activity of each neuron saved to file. Afterwards, each cell is modeled with biophysically detailed multicompartment morphologies and the stored spikes of all the presynaptic neurons are used as activation times for synaptic input onto these neurons in a simulation where the extracellular potentials are calculated [Hagen et al., 2016; Senk et al., 2018].

We used the large-scale point-neuron cortical microcircuit model from Potjans and Diesmann [2014]; Hagen et al. [2016], which has ~80 000 neurons divided into 8 different cortical populations, one excitatory and one inhibitory, across four layers (L2/3 - L6), and can exhibit a diverse set of spiking dynamics including different oscillations and asynchroneous irregular network states [Hagen et al., 2016; Brunel, 2000]. We chose the scenario with transient thalamocortical input, and the only difference from the original simulation by Hagen et al. [2016] was the added calculation of current dipole moments and EEG signals. We simulated transient thalamic synaptic input to layers 4 and 6 (Fig. 5**A**), and after the spikes had been mapped onto the multicompartment cell models (Fig. 5**B**), we calculated the LFP (Fig. 5**C**) similarly to Hagen et al. [2016] (their Fig. 1), in addition to the current dipole moments of each cell.

For all cell populations, we found that the current dipole moments from individual cells could show large transient responses to thalamic input (Fig. 5**D**; gray lines show current dipole moment from individual cells in two example populations: L5 inhibitory (L5I) and L5 exitatory (L5E)), but for all inhibitory populations the thalamic response was not visible in the average current dipole moment (Fig. 5**D**; black lines, L5I). The same was true for the current dipole moment components perpendicular to the depth axis for excitatory populations (Fig. 5**D**; L5E, *p_x_*, *p_y_*, black lines), but not for the component along the depth axis which had a substantial average response to the thalamic input (Fig. 5**D**; L5E, *p_z_*, black line). These observations imply, as previously noted, that only the *z*-component of the current dipole moment from excitatory populations can be expected to contribute significantly to the EEG signal.

Our findings invite a simplified approach to calculate the EEG signal: Instead of calculating all single-cell EEG contributions and summing them (taking into account the position of the individual cells, similarly to what is done for the LFP signal), we can compute a single summed *p_z_*-component from all neurons in each pyramidal cell population, place it in the center of the population column, and calculate the resulting simplified EEG signal. This approximation can be expected to be reasonable when the population radius is small compared to the distance from the population center to the EEG electrode. Note that the distance from the top of cortex to the top of the head is typically ~10 mm, while the radius of the present simulated population is only ~0.5 mm (Fig. 5; population outline in **B** is drawn in red in **E**). Indeed, when we combined the current dipole moments with the four-sphere head model (Fig. **5E**), we found that the full EEG signal that was calculated as the sum of the EEG contribution from each of the ~80 000 cells at their respective positions, was in fact indistinguishable from the simplified EEG signal (Fig. **5F**). This implies that the EEG signal from the simulated cortical activity can be fully represented by a single time-dependent variable for each pyramidal cell population.

We also compared the relative amplitude of the EEG signal from each population, and found that for the present example, the excitatory population of L2/3 was the dominant source of the EEG signal (Fig. **5F**). Note, however, that we expect this observation to be somewhat model-dependent, and that strong general claims about the contribution of different pyramidal cell populations to the EEG signal cannot be made from this example study alone.

### 3.6. Dipole approximation in complex head models

Even though the four-sphere head model is convenient for generic EEG studies, many applications such as accurate EEG source analysis, may require more detailed head models [Dale et al., 1999; Vorwerk et al., 2014]. The construction of such complex head models is dependent on expensive equipment, that is magnetic resonance imaging (MRI), to map the electrical conductivity of the entire head at resolutions of ~0.5-1.0 mm^3^ [Huang and Parra, 2015; Huang et al., 2016]. Afterwards, numerical techniques such as the Finite Element Method (FEM) [Logg et al., 2012] can be used to calculate the signal at the EEG electrodes for arbitrary arrangements of current dipoles in the brain, but at a high computational cost. The number of computing hours is, however, reduced by applying the reciprocity principle of Helmholtz. The reciprocity principle states, in short, that switching the location of a current source and a recording electrode will not affect the measured potential [Malmivuo and Plonsey, 1995; Ziegler et al., 2014; Huang et al., 2016; Dmochowski et al., 2017]. This implies that it suffices to use FEM to calculate the lead field in the brain from virtual current dipoles placed at each of the EEG electrodes. From the lead field matrix, we can infer the potential at the EEG electrodes, given an arbitrary arrangement of current dipoles in the brain. Luckily, several such pre-solved complex head models are freely available, and one example is the *New York Head* (NYH) (Fig. **6A**), which we have applied here (Huang et al. [2016]; parralab.org/nyhead/).

To illustrate the use of pre-solved complex head-models, we inserted the current dipole moment obtained from the cortical column model in section 3.5 into the New York Head model (Fig. **6A**), at two manually chosen positions: one in the parietal lobe, and one in the occipital lobe, see stars in Fig. **6C,E**, respectively. In both cases, the current dipole moment was oriented along the normal vector of the brain surface, and the EEG signal was calculated. For comparison with a simplified head model, we inserted the same current dipole moment into the four-sphere head model (Fig. **6B**) at locations comparable to the dipole positions chosen in the occipital and parietal lobe in the NYH model: the locations in the four-sphere model were chosen close to the brain surface, such that the distance from dipole position to the closest electrode (Fig. 6**D, F**, stars) and the brain surface normal vectors were similar to the respective positions in the NYH model.

The computed EEG signals from the two head models were in this case relatively comparable in both spatial shape and amplitude (Fig. 6**C-F**). The two head models also generated EEG signals of the same temporal shape, as expected, but while the four-sphere head model gave very similar EEG amplitudes for the two different dipole locations, the EEG amplitudes from the complex head model was much more variable, even for similar distances to the closest electrode (Fig. 6**G**, **H**).

The higher variability of the complex head model was also apparent in the decay of the maximum EEG amplitude with distance, which was perfectly smooth, exponential-like [Nunez and Srinivasan, 2006], and very similar for the two locations in the four-sphere model, but very variable for the complex head model, although with the same general shape (Fig. 6**I**, **J**).

Note that despite the complexity, the NYH model is substantially faster than the four-sphere model. In order to simulate the EEGs from a dipole moment vector containing 1200 timesteps, the NYH model execution times were ~ 0.4 s, while the four-sphere model needed ~ 1.5 s.

## 4. Discussion

### 4.1. Summary

In this paper, we have introduced an approach for reducing arbitrary simulated neural activity from biophysically detailed neuron models to single current dipoles (Fig. 1). We verified that the approach was applicable for calculating EEG, but generally not for ECoG signals (Fig. 2-3), and gave examples of how reducing neural activity to a single dipole can be a powerful tool for investigating and understanding single-cell EEG contributions (Fig. 3-4). Furthermore, we demonstrated that the presented approach could easily be integrated with existing large-scale simulations of neural activity. Moreover, we showed how single dipoles are useful for constructing compact representations of the EEG contributions from entire neural populations, with methods still firmly grounded in the underlying biophysics (Fig. 5). Finally, we demonstrated how the simulated current dipoles, from single cells or large neural populations, can be directly inserted into complex head models for calculating more realistic EEG signals (Fig. 6).

### 4.2. Application of current dipoles for computing EEG, MEG and ECoG signals

We have highlighted that the calculation of current dipoles from neural activity is cleanly separated from the calculation of the ensuing EEG signals. Since MEG sensors like EEG electrodes are positioned far away from the neural sources, the same is true for MEG signals. The calculated current dipoles can therefore also be used in combination with simplified or detailed frameworks for calculation of MEG signals, for example by following methods outlined in Hagen et al. [2018]; Ilmoniemi and Sarvas [2019].

ECoG electrodes are in general positioned closer to the neural sources. For our example simulations of the ECoG signal generated by individual neurons, we found that use of the single-dipole approximation gave substantial errors (Fig. 2). Thus for computation of ECoG signals, the standard compartment-based formalism or the multi-dipole approach (Fig. 1) requiring much more computational resources, may be required. Here an alternative to using full head models is to use the method of images, taking into account the discontinuity of electrical conductivity at the cortical surface [Pettersen et al., 2006; Hagen et al., 2018]. Note that while we here used LFPy 2.0 [Hagen et al., 2018, 2019], a python interface to NEURON [Carnevale and Hines, 2006], calculation of current dipole moments can easily be implemented into any framework where the transmembrane currents are available, through the simple formula given in eq. (7).

### 4.3. Generalization to non-compartmental models

EEG and MEG recordings reflect neural activity at the s ystems-level [Pesaran et al., 2018; Einevoll et al., 2019]. Here, we have focused on calculating current dipoles from detailed multi-compartment neuron models, but neural modeling at the systems-level is often based on higher levels of abstraction, like point neurons [Linssen et al., 2018] or firing rate populations [Sanz-Leon et al., 2013]. Calculation of electric or magnetic signals from such higher-level neural simulations must in general rely on some kind of approximation trick, since neurons require a spatial structure to be capable of producing electromagnetic signals [Einevoll et al., 2013a]. One such trick that we took advantage of here is the hybrid scheme [Hagen et al., 2016]. This two-step scheme involves neural network activity first being simulated by point neurons, before the resulting spike trains are replayed onto multi-compartment neuron models for calculating LFP and EEG signals (sec. 3.5; Fig. 5).

Further, the hybrid scheme can be generalized to also allow for calculation of EEG/MEG signals from firing-rate models by using the so-called kernel method, which has previously been successfully applied to the LFP [Hagen et al., 2016; Skaar et al., 2020; Teleńczuk et al., 2020]. In practice, this can be done in two steps: First, simultaneously activating all outgoing synapses from a specific (presynaptic) simulated population, and recording the total current dipole moment of the response (the kernel) [Hagen et al., 2016]. Second, computing the EEG/MEG contribution stemming from this (presynaptic) population by convolving the kernel with the population firing rate, and applying an appropriate forward model. Here, the firing rate would be obtained separately in point neuron network models or firing-rate models. In this way, the basic biophysics of EEG and MEG signals from synaptic activation of multi-compartment neuron models is included, avoiding, however, computationally heavy multicompartmental modeling of spiking dynamics. The calculated current-dipole kernels should be applicable for different kinds of input to the original network model, but would in general have to be recomputed for changes to cell or synaptic parameters.

### 4.4. Connection to other work

Calculation of current dipole moments from morphologically complex cell models has been pursued before, for example to study the EEG and MEG contribution of spiking single cells [Murakami and Okada, 2006], or to study how the synaptic input location affects the current dipole [Lindén et al., 2010; Ahlfors and Wreh II, 2015]. Important work on EEG interpretation in terms of the underlying neural activity has also previously been done through use of “minimally sufficient” biophysical models, see for example Murakami et al. [2002, 2003]; Jones et al. [2007, 2009]; Sliva et al. [2018]; Neymotin et al. [2020]. Here, “minimally sufficient” means that the cell models only had minimally needed multi-compartment spatial structure (point neurons cannot produce current dipole moments), only considered a few cell types, and employed simple synaptic connection rules. In particular, the Human Neocortical Neurosolver (HNN) [Neymotin et al., 2020] enables researchers to link measured EEG or MEG recordings to neural activity through a pre-defined canonical neocortical column template network. HNN comes with an interactive GUI, designed for users with little or no experience in computational modeling, and might therefore be an appropriate choice for researchers seeking to gain a better understanding of their EEG/MEG data. However, while the use of such minimally sufficient models allows for quick and direct comparison between simulated and recorded EEG signals, it is not (presently) compatible with simulating EEG or MEGs from biophysically detailed single cell- or network models, constructed from detailed experimental data [Reimann et al., 2013; Egger et al., 2014; Markram et al., 2015; Hagen et al., 2016; Gratiy et al., 2018; Arkhipov et al., 2018; Billeh et al., 2020].

A more high-level approach for simulating MEG/EEG signals from the underlying neural activity has been pursued through neural field or neural mass models [Jirsa et al., 2002; David and Friston, 2003; Coombes, 2006; Deco et al., 2008; Bojak et al., 2010; Ritter et al., 2013], which aim to model the evolution of coarse-grained variables such as the mean membrane potential or the firing rate of neuron populations. Such coarse-graining drastically reduces the number of parameters and the computational burden of the simulation, and can be used to study the interplay among entire brain regions, and indeed run whole-brain simulations. The Virtual Brain (TVB) is an excellent example of a software for whole-brain network simulations [Sanz-Leon et al., 2013, 2015; Ritter et al., 2013], where detailed and potentially personalized head models can be combined with tractography-based methods identifying the connectivity between brain regions [Sanz-Leon et al., 2013]. To calculate measurement modalities like MEG and/or EEG signals from neural field or neural mass models, it is typically assumed that the population current dipole moments are roughly proportional to, for example, the average excitatory membrane potential [Bojak et al., 2010; Ritter et al., 2013]. Further, EEGs can be calculated from the resulting current dipole moments in combination with head models as presented in this paper, or through other softwares or techniques [Gramfort et al., 2014]. This suggests an intriguing future development, where one could apply the above-mentioned kernel method based on biophysically detailed neuron models to substantially increase the accuracy of LFP, EEG and MEG predictions from high-level large-scale simulations of neural activity.

### 4.5. Outlook

EEG and, later, MEG signals have been an important part of neuroscience for a long time, but still very little is known about the neural origin of the signals [Cohen, 2017]. A better understanding of these signals could lead to important discoveries about how the brain works [Lopes da Silva, 2013; Uhlirova et al., 2016; Pesaran et al., 2018; Ilmoniemi and Sarvas, 2019], and provide new insights into mental disorders [Mäki-Marttunen et al., 2019a; Sahin et al., 2019]. This work lays some of the foundation for obtaining a better understanding of EEG/MEG recordings, by allowing easy calculation of the signals from arbitrary neural activity. The presented formalism is well suited for modeling EEG/MEG contributions from various potential neural origins, including different cell types, different ion channels and different synaptic pathways. For example, to study the effect of calcium spikes [Suzuki and Larkum, 2017], I_h_ currents [Ness et al., 2016, 2018; Kalmbach et al., 2018], or gene expression on EEG signals [Mäki-Marttunen et al., 2019b], one only needs to know how the *z*-component of the resulting population current dipole is affected. This decoupling of the current dipole moment and head model allows for easier investigation and improved understanding of the origin of the EEG/MEG signal.

EEG/MEG measurements are often used for source localization, aiming to identify the underlying cortical current dipoles [Nunez and Srinivasan, 2006; Gramfort et al., 2014; Ilmoniemi and Sarvas, 2019]. However, such reconstructed current dipoles are generic in the sense that they are typically not intended to represent specific neural populations. By allowing for calculation of current dipoles from cortical populations, the work presented here takes a step towards consolidating the, so far, mostly separate scientific disciplines of neural modeling and EEG/MEG data analysis (but see also Neymotin et al. [2020]).

While there are many examples of detailed biophysical modeling of neural activity improving interpretation of measured intracranial extracellular potentials in lab animals [Einevoll et al., 2007; Blomquist et al., 2009; McColgan et al., 2017; Luo et al., 2018; Chatzikalymniou and Skinner, 2018; Teleńczuk et al., 2019], much less has been done for human EEG/MEG signals. This is natural given that studies of healthy human brains necessarily are limited to non-invasive technologies [Lopes da Silva, 2013; Uhlirova et al., 2016; Cohen, 2017]. However, given all the valuable insights that could be gained from an increased understanding of non-invasive measurements of neural activity in humans, an important challenge in modern neuroscience is to build on the mechanistic insights from animal studies and use them for understanding non-invasive signals in humans [Lopes da Silva, 2013; Uhlirova et al., 2016; Cohen, 2017; Einevoll et al., 2019; Mäki-Marttunen et al., 2019a]. The approach for calculating EEG/MEG signals in this paper should therefore ideally be used in combination with animal studies simultaneously measuring multisite laminar LFP (and MUA) signals within cortex, as well as EEG/MEG signals (see for example Bruyns-Haylett et al. [2017]) [Cohen, 2017].

Today, we have a reasonably good understanding of how single neurons operate, that is, how they respond to synaptic input, and how multitudes of synaptic inputs combine to produce action potentials [Einevoll et al., 2019]. Similarly, we can, to a high degree, explain the measurement physics of EEG/MEG, that is, how neural currents affect electromagnetic brain signals recorded outside of the head [Nunez and Srinivasan, 2006; Cohen, 2017; Ilmoniemi and Sarvas, 2019]. The challenge of understanding EEG/MEG signals is therefore closely related to the greatest challenge in modern neuroscience: understanding neural networks. Making sense of such complicated dynamical systems typically requires computational modeling [Einevoll et al., 2019], but the complexity of neurons, and the complexity and size of the neural networks involved in even the simplest of cognitive tasks, makes this a daunting challenge. The steady increase in available computing power, in combination with the ever-increasing knowledge on synaptic connectivity patterns is, however, making this approach more and more attractive [Reimann et al., 2013; Egger et al., 2014; Markram et al., 2015; Hagen et al., 2016; Gratiy et al., 2018; Arkhipov et al., 2018; Reimann et al., 2019; Billeh et al., 2020]: Today, there are several ongoing research projects pursuing such modeling efforts, for example at the Allen Institute for Brain Science and in the Human Brain Project [Einevoll et al., 2019]. While biophysically detailed, large-scale neural simulations are still in their infancy, we expect these simulations to become an increasingly important research tool in neuroscience [Einevoll et al., 2019]. The presently described method enables EEG/MEG simulations combining detailed neural simulations with realistic head models. We believe that this approach will help shedding light on the neural origin of EEG/MEG signals, and help us take full advantage of these important brain signals in the future.

## Acknowledgements

This work received funding from the European Union Horizon 2020 Research and Innovation Programme under Grant Agreement No. 785907 and No. 945539 [Human Brain Project (HBP) SGA2 and SGA3], the Norwegian Ministry of Education and Research through the SUURPh Programme and the Norwegian Research Council (NFR) through COBRA (No. 250128), NOTUR (No. NN4661K) and DigiBrain (No: 248828).

## Notes

### Competing Interest Statement

The authors have declared no competing interest.

